# Phenotypic and genomic changes in enteric *Klebsiella* populations during long-term ICU patient hospitalization: The role of RamR regulation

**DOI:** 10.1101/2024.04.30.591843

**Authors:** Benedicte Langlois, Francois Guerin, Christophe Isnard, Clement Gakuba, Damien Du Cheyron, Jean-Christophe Giard, Sylvain Brisse, Simon Le Hello, Francois Gravey

**Affiliations:** Univ de Caen Normandie, Univ Rouen Normandie, INSERM, DYNAMICURE UMR1311, Caen, France; CHU Caen, Department of Infectious Agents, Bacteriology, Caen, France; CHU Caen, Department of Surgical Intensive Care, Caen, France; CHU Caen, Department of Medical Intensive Care, Caen, France; Institut Pasteur, Université Paris Cité, Biodiversity and Epidemiology of Bacterial Pathogens, Paris, France

## Abstract

Acquired antimicrobial resistance and metabolic changes are central for bacterial host adaptation during the long-term hospitalization of patients. We aimed to analyze the genomic and phenotypic evolution of enteric *Klebsiella* populations in long-term intensive care unit (ICU) patients.

Weekly rectal swabs were prospectively collected from all patients admitted to the ICU in a teaching hospital from December 2018 to February 2019. The inclusion criterion for patients was hospitalization for more than 15 days in the ICU without any history of hospitalization or antibiotic treatment for the three months prior to admission. Among them, enteric *Klebsiella pneumoniae* species complex (KpSC) populations were detected. For each isolate, extensive antimicrobial resistance profiles were determined using the disk diffusion method, and the whole genome was sequenced using an Illumina platform. *In silico* typing methods, such as multilocus sequence typing (MLST), core-genome MLST, SNP typing, resistome characterization and mutation point detection, were applied.

During the study period, 471 patients were admitted to ICUs. Among them, 21 patients met the inclusion criteria, and only five patients (24%) carried unique and distinct KpSC populations during two to ten weeks in the gut that as detected at admission and excluding acquisition during the ICU stay. One patient showed a rare ST1563 *K. variicola* persistent carriage for seven consecutive weeks, which displayed important antimicrobial resistance phenotype changes in the two last weeks. In-depth *in silico* characterization and RNA sequencing of these strains revealed a mutation within the *ramR* transcriptional regulator resulting in overexpression of the *ramA* regulator and decreased expression of *acrR*. These modifications are implicated in multidrug resistance, biliary salt tolerance and other bacterial functions.

This study revealed the importance of endogenous colonization of KpSC populations in the gut throughout the patient’s long-term ICU stay and highlighted the role of *ramR* regulation in microbial adaptation.

**Author summary:** *Klebsiella variicola* belongs to a large bacterial complex named the *Klebsiella pneumoniae* species complex (KpSC). These bacteria are largely involved in nosocomial infection and are able to colonize human gut microbiota during hospitalization and/or develop antimicrobial resistance during the hospital stay. In this work, we aimed to determine the prevalence at admission and adaptation of persistent KpSC populations in the gut of long-term ICU patients. Among 471 patients admitted, 21 were hospitalized for more than 15 days, and 5 carried a unique and distinct endogenous KpSC. *K. variicola* was detected in two of the five patients, and antibiotic resistance was detected during long-term hospitalization in these patients. One *K. variicola* strain became cross-resistant to chloramphenicol, quinolones and tigecycline after the seventh week of hospitalization. *In silico* analyses revealed the persistence of a rare ST 1563 *K. variicola* population with a mutation in the *ramR* transcriptional regulator, which controls RND efflux pump expression and antibiotic efflux. This mutation also impacts tolerance to biliary salts and probably biofilm formation.

In conclusion, a mutation in an important transcriptional regulator, *ramR,* could be involved in not only antimicrobial resistance but also facilitate persistent *K. variicola* colonization.

## Introduction

The *Klebsiella pneumoniae* species complex (KpSC) is a wide complex belonging to the *Enterobacterales* order and comprises seven phylogroups identified via phylogenetic analyses, including *K. pneumoniae*, *K. variicola* and *K. quasipneumoniae* [1,2]. They are naturally present in soil, water and plants, as well as in the digestive tract of animals [3]. They constitute a part of the human digestive microbiota but are inconsistently found among rectal sample cultures. Indeed, the digestive carriage frequencies are estimated, depending on the selective detection method, to be between 6% and 35% in the general population. This frequency increases during hospitalization to 77%, notably among patients hospitalized in intensive care units (ICUs) [4–8]. Studies have identified risk factors for intestinal colonization by KpSC populations regardless of the antimicrobial susceptibility profile, such as the length of hospital stay, the use of nonantibiotic molecules (proton pump inhibitors and nonsteroid anti-inflammatory treatments) and antibiotic pressure, particularly for populations that produce extended-spectrum β-lactamases (ESBLs) or carbapenemases [7,9,10]. Furthermore, gastro-internal carriage is associated with a greater risk of KpSC infections, such as urinary and respiratory tract infections, bacteraemia, and septic shock [5,11]. The risk of deep infection ranges from 5.2% to 16% for patients with digestive KpSC carriage versus 1.3% to 3% for people without digestive KpSC carriage, regardless of the antibiotic resistance profile.

The KpSC includes bacteria with important genomic plasticity that confer adaptability to persist and resist the environment [5,12]. Regarding digestive colonization, some genes could be implicated in colonization and perhaps in persistence, notably those involved in sugar metabolism that possibly play a role in membrane stability and lipopolysaccharide synthesis (*galET* operon, *waa* operon), genes linked to nitrogen metabolism (*ntrC*), and genes involved in bacterial growth and the regulation of aerobe-anaerobe metabolism (*arcA*, *arcB*) [13–15]. Modulation of these bacterial pathways promotes KpSC gut persistence, which could allow bacteria to adapt to the hospital environment during long-term stays. Bacterial adaptation could develop biocide resistance, with environment and human colonization leading to nosocomial infections and/or important outbreaks in ICUs [5,16]. Even if there are multiple factors implicated in the adaptation of the KpSC carried by long-term ICU patients, the digestive tract seems to be a suitable site for antimicrobial resistance. Thus, specific ESBL-KpSC and carbapenemase-producing KpSC (CP-KpSC) clones are common causes of healthcare-associated infections leading to therapeutic failure and increased morbi-mortality [17–19]. However, the acquisition of encoding genes is not the only way to develop multidrug resistance (MDR) phenotypes in these species. Indeed, efflux pump dysregulation, including the upregulation of resistance-nodulation-division (RDN) pumps, such as the AcrAB or OqxAB pumps, which increase minimal inhibitor concentrations (MICs) toward tigecycline and quinolones, is also largely involved [20–22]. Worryingly, evolutionary convergence of MDR and hypervirulent *K. pneumoniae* populations has recently emerged and has posed serious therapeutic challenges [23].

In the present 2-month study, we aimed to estimate and describe the long-term frequency of KpSC carriage in ICU patients. Extensive phenotypic and genomic analyses were performed on persistent KpSC populations identified in the gut of long-term hospitalized patients to (i) describe the dynamics of digestive carriage and (ii) define the mechanisms of antimicrobial resistance (AMR) acquisition and metabolic pathway modifications. An in-depth molecular investigation was performed for a *K. variicola* isolate displaying an important change in the multidrug resistance phenotype and metabolic pathway from the seventh week of carriage.

## Results

### Epidemiology

A total of 471 patients were admitted to ICUs between December 2018 and February 2019 at Caen Normandy University Hospital. Among them, only 21 patients (4.5%) met the inclusion criteria as follows: hospitalization for more than 15 days in the ICU and had not been hospitalized or had not received antibiotic treatment for the three months prior to admission. The median length of stay for long-term stays was 33 days [min: 17; max: 129], with a median number of rectal samples of five [min: 3; max: 13] (Fig 1). These patients were admitted for sudden events such as hemorrhagic or thrombotic stroke (n=10), trauma (n=6), complicated flu (n=1), digestive hemorrhage (n=1), septic shock secondary to peritonitis (n=1), acute coronary syndrome (n=1) or pulmonary embolism (n=1). Among these 21 long-stay patients, both ZKIR qPCR (with a median CT of 16.7 [min: 11.4; max: 33.2]) and culture were positive for eight patients, which indicates a prevalence of enteric KpSC carriage of 38% (8/21) among long-term stay patients in our ICUs. Among them, five patients (24%), aged between 31 and 76 years, three women and two men, were continually colonized by KpSC strains for two to ten weeks. Species identification revealed that three patients were colonized by *K. pneumoniae* and two patients were colonized by *K. variicola*. Four patients were infected with KpSC species throughout their hospitalization, and one experienced KpSC clearance after four weeks of colonization without concomitant antibiotic treatment (S1 and S2 Appendix). The characteristics of the bacterial isolates from the five patients are available in S3 Appendix. No long-term carrier patients developed deep KpSC infection.

### Multilocus sequence typing (MLST) and single nucleotide polymorphism (SNP) typing of the KpSC isolates from the five long-term patients

#### MLST

KpSC isolates from the five patients showed they belonged to a unique and distinct ST each: ST36, ST163, ST1563, ST1791 and ST405. Except for the last one, these STs were not related to the KpSC populations already described in our local genomic database (n= 236 KpSC populations sequenced between 2015 and 2019 according to our routine surveillance in the ICU wards in our hospital) [24]. The last population was a *K. pneumoniae* ST405 ESBL-producing population, which showed colonization in the patient from the second week of hospitalization until the end of hospitalization. This strain was prevalent in our database (n=51/236, 21.4%) [24].

**Fig 1.**
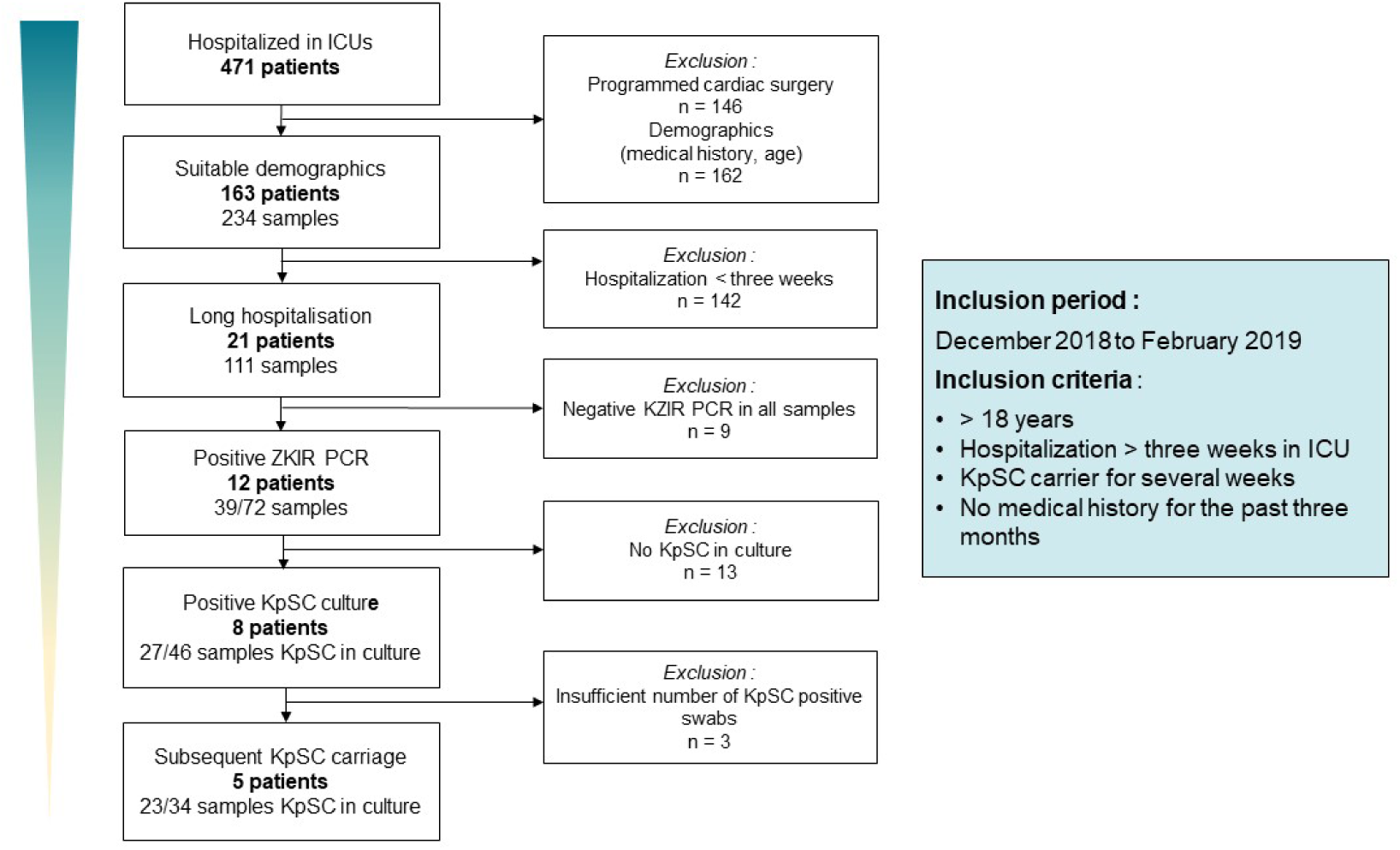
Study Flow Chart of Patient Inclusion from ICUs at Caen University Hospital Center. KpSC: *K. pneumoniae* species complex. n: number of patients.

#### Core SNP analyses

A pairwise comparison of isolates from the five KpSC populations revealed relative genome stability between all the isolates of each unique patient (8 ST36 isolates: median of 6 SNPs [0;12]; 3 ST163 isolates: median of 6 SNPs [6;6]; 8 ST1563 isolates: median of 1 SNP [0;2]; 3 ST1791 isolates: median of 4 SNPs [3;7]; and 3 ST405 isolates: median of 2 SNPs [0;4]). The remaining ESBL-producing strain was distant from our local epidemic cluster ST405 strains, with more than 2000 SNPs [24].

### Biocide susceptibility screening, clinical case report and genomic analyses

Among these five patients, only the two *K. variicola* populations displayed AMR profile changes during long-term hospitalization: ST1791 and ST1563.

For the *K. variicola* ST1791 isolates from the second round of swabs, which were collected 16 days after the patient was admitted to the ICU, there was a reduction in susceptibility diameters for macrolides (azithromycin [10 mm]), aminoglycosides (streptomycin [12 mm]) and β-lactams (mecillinam [8 mm]) compared to the isolates collected during the first week after the patient was admitted to the ICU (S4 Appendix). This resistance was concomitant with the appearance of genes coding for resistance to macrolides (*mphA)*, aminoglycosides (*aadA5* and *strAB)* and a β-lactamase (*bla_TEM1D_*) carried on a putative IncFIB plasmid (S3 Appendix). Isolates sampled during the fourth round of swabs, which were collected after one month of hospitalization, showed β-lactam diameter reduction (ceftazidime [13 mm], cefepime [7 mm], and aztreonam [11 mm]) that could be attributed to a mutation in *ompK35* porin (S3 Appendix). KpSC populations were not detected after the fourth week of ICU hospitalization. Because of the absence of persistence, these patient isolates were not further explored.

For the ST1563 digestive *K. variicola* isolates collected during the first six weeks of hospitalization, a natural antimicrobial resistance profile (i.e., resistance to ampicillin and ticarcillin), as defined by the European Committee of Antimicrobial Susceptibility Testing (EUCAST), was observed. Modification of the AMR profile also appeared in the isolates collected during the seventh week after admission (ESSO7) and persisted through the eighth week (ESSO8). Several molecules were impacted, with diameter restrictions ranging from 5 mm to 11 mm: quinolones (nalidixic acid [7 mm], ciprofloxacin [5 mm], norfloxacin [6 mm] and levofloxacin [6 mm]), chloramphenicol [11 mm], tigecycline [5 mm], some β-lactams (temocillin [6 mm], and cefoxitin [7 mm]) (S4 Appendix). Loss of susceptibility was confirmed by minimum inhibitory concentrations (MICs) in microdilutions, with reductions of 8-fold for tigecycline, chloramphenicol and cefoxitin and 4-fold for ciprofloxacin, nalidixic acid and temocillin (Supplementary S5). Interestingly, the strains isolated during the seventh sampling time point presented two different AMR profiles: one isolate with a natural AMR profile and four isolates with a new AMR profile, whereas all 5 isolates collected during the eighth sampling time point were resistant to the abovementioned molecules. To investigate further, microdilutions were made for the three antiseptics frequently used in hospitals and no variation in susceptibility was observed. The MIC of povidone-iodine was 1563 mg/L for all isolates, the MIC of didecyldimethylammonium chloride (DDAC) ranged between <1 mg/L and 4 mg/L and the MIC of chlorhexidine ranged between 4 and 32 mg/L. Thus, no tolerance to the antiseptics tested was observed in the five KpSC populations, and the profiles remained stable over the entire time period (S5 Appendix).

In terms of the patient from which the *K. variicola* ST1563 strain was isolated from, she was a 61-year-old woman who was suddenly admitted to the ICU in January 2019 following a stroke. She was hospitalized for eight weeks, had no medical history or comorbidities before her stay and was discharged to the neurosurgery unit for the postacute phase. During her ICU stay, eight rectal swabs were collected (one per week). No MDR bacteria were found, but the KpSC screening PCR was positive for all samples, and *K. variicola* colonies were isolated during the last seven weeks of her stay. Regarding antibiotic treatments, the patient received three days of broad-spectrum antibiotics with piperacillin-tazobactam relayed by oxacillin for seven days due to *S. aureus* pneumopathy for a week during the second week of hospitalization. Moreover, the patient received erythromycin twice at a dose of 250 mg four times a day for six days to stimulate digestive motility (Fig 2).

**Fig 2.**
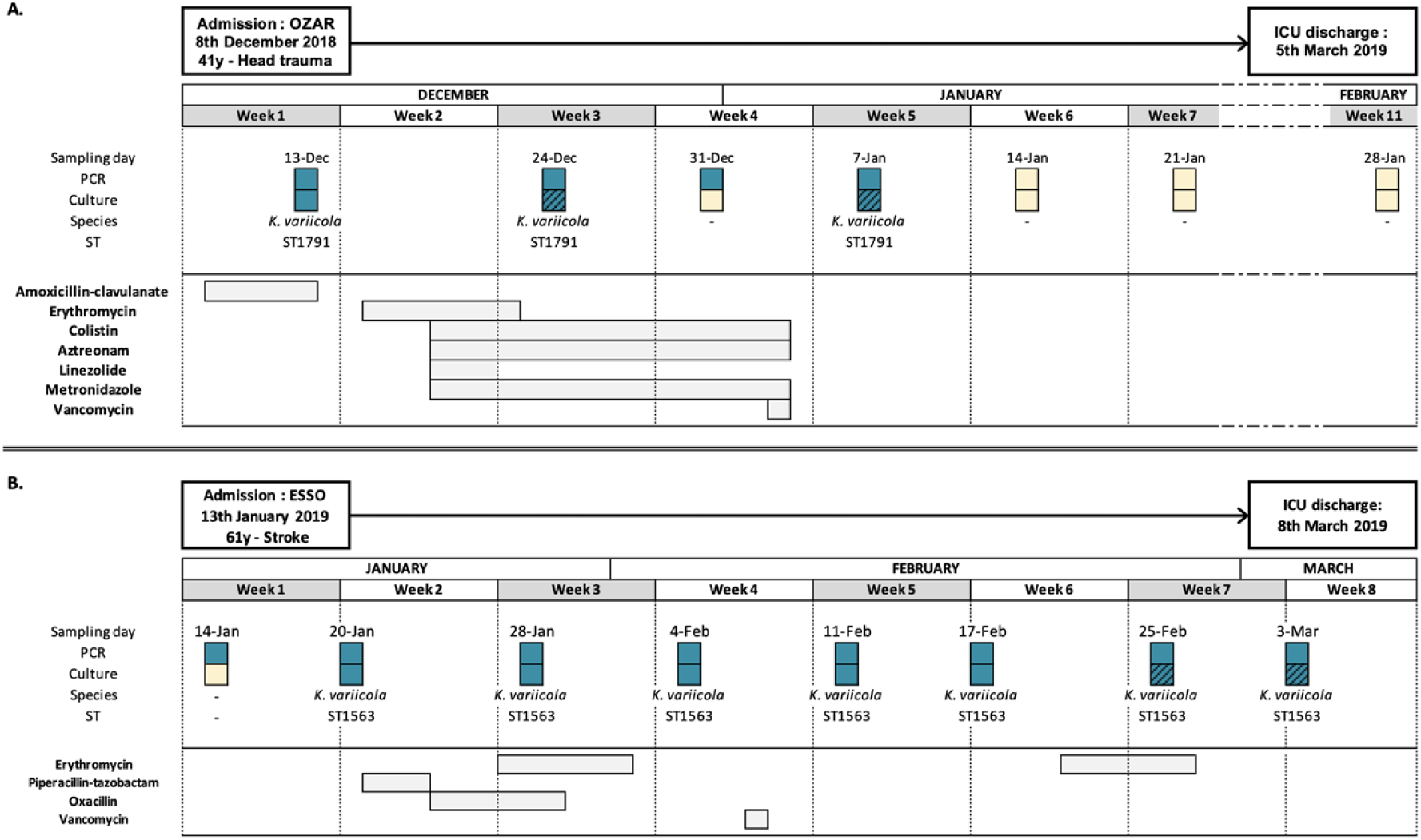
Patient Hospitalization Timelines. (A) OZAR patient, (B) ESSO patient. Blue: positive; yellow: negative; dashed: decreased antibiotic susceptibility.

### Genomic/phenotypic analyses of the *K. variicola* ST1563 isolates

One isolate was sequenced for each sampling time point, except for the seventh sampling point, where two AMR profiles were observed, and thus both phenotype strains were sequenced. For all eight isolates, WGS data revealed the presence of the plasmid genes *tetD* and *bla_SHV1_*and the chromosomal resistance gene *bla_LEN17_*. The resistome was stable from the second to the eighth week, indicating that no acquired resistance gene could explain the AMR profile switch observed during the seventh week (S3 Appendix). In fact, the AMR phenotype evolution observed in ESSO7 and ESSO8 was concomitant with a single genomic mutation (c.454 G>T) in the *ramR* transcriptional regulator. This unique *ramR* nonsilent mutation leads to a RamR protein modification, swapping an aspartate to a tyrosine at position 152.

To validate the involvement of RamR in the loss of antibiotic susceptibility, ESSO7 and ESSO8 were complemented with a wild-type sequence of *ramR* extracted from the ESSO2 strain isolated at the second sampling time point: ESSO7Ω*ramR* and ESSO8Ω*ramR*. Strains with a mutated *ramR* gene presented a shorter doubling time and a greater growth rate in the presence of erythromycin (256 mg/L) or biliary salts at a high concentration (40 mg/mL) than other isolates with nonmutated *ramR* (S6 and S7 Appendix). However, no difference in growth was observed under the other conditions tested, including acidic pH, starvation, H_2_O_2_, 22 °C temperature and hyperosmotic stress (data not shown). Strains complemented with a wild-type *ramR* (ESSO7Ω*ramR* and ESSO8Ω*ramR*) had the same growth profile as the nonmutated *ramR* strains, suggesting that the *ramR* mutation is involved in digestive adaptation.

Next, the MICs of chloramphenicol, ciprofloxacin and tigecycline in the presence of phenylalanine-arginine beta-naphthylamide (PAβN) were found to be significantly decreased, reaching values similar to those of the wild-type strains (≥4-fold reduction) (Table 1). For temocillin, the reduced MIC under inhibitor pressure was not as pronounced as that of the other molecules (2-fold reduction). This test confirmed the involvement of these RND pumps in the AMR profile observed in the *K. variicola* ST1563 strains. Furthermore, the MICs of ESSO7Ω*ramR* and ESSO8Ω*ramR* were identical to those of the wild-type strains. These observations support the hypothesis that this single mutation in *ramR* is responsible for the reduced susceptibility to antimicrobial agents and the digestive environment.

**Table 1.** MICs of ESSO for Temocillin, Chloramphenicol, Ciprofloxacin and Tigecycline. PAbN was used at a concentration of 20 mg/L.

| Strains | MIC (mg/L) |  |  |  |
| --- | --- | --- | --- | --- |
|  | TEM | CHL | CIP | TGC |
| ESSO2 | 4 | 8 | 0.03 | 0.5 |
| ESSO3 | 8 | 8 | 0.03 | 0.5 |
| ESSO4 | 4 | 8 | 0.03 | 0.5 |
| ESSO5 | 4 | 8 | 0.03 | 0.5 |
| ESSO6 | 4 | 8 | 0.03 | 0.5 |
| ESSO7 | 16 (4) | 64 (8) | 0.125 (4) | 4 (8) |
| ESSO8 | 16 (4) | 64 (8) | 0.125 (4) | 2 (4) |
| ESSO7 + PAβN | 8 (-2) | 0.5 (-128) | 0.03 (-4) | 0.5 (-8) |
| ESSO8 + PAβN | 8 (-2) | 0.5 (-128) | 0.03 (-4) | 0.25 (-8) |
| ESSO7ΩpBAD202_*ramR* | 2 (-8) | 2 (-32) | 0.03 (-4) | 0.25 (-8) |
| ESSO8ΩpBAD202_*ramR* | 4 (-4) | 2 (-32) | 0.015 (-8) | 0.25 (-8) |
| ESSO7ΩpBAD202_empty | 8 | 32 | 0.125 | 4 |
| ESSO8ΩpBAD202_empty | 16 | 32 | 0.25 | 4 |

CHL: chloramphenicol, CIP: ciprofloxacin, MIC: minimal inhibitory concentration; TEM: temocillin, TGC: tigecycline

### Transcriptomic levels of the *ramAR* locus and *acrAB*-*tolC* genes

To validate the impact of the *ramR* mutation on RND pump regulation, quantitative reverse transcription PCR of the *ramAR* locus, *romA* gene and *acrAB*/*tolC* RND pump was performed. The calculated fold change (FC) highlighted significant overexpression of *ramA* (FC: 20), *romA* (FC: 21) and *ramR* (FC: 6) in the group with mutated *ramR,* which confirmed the loss of repression of the RamR protein on the *ramA* and *romA* genes. In strains overexpressing RamA, the RND pump directly controlled by RamA showed nonsignificant overexpression of *acrB* (FC: 1.78), *acrA* (FC: 1.55) and *tolC* (FC: 1.48) (Fig 3).

**Fig 3.**
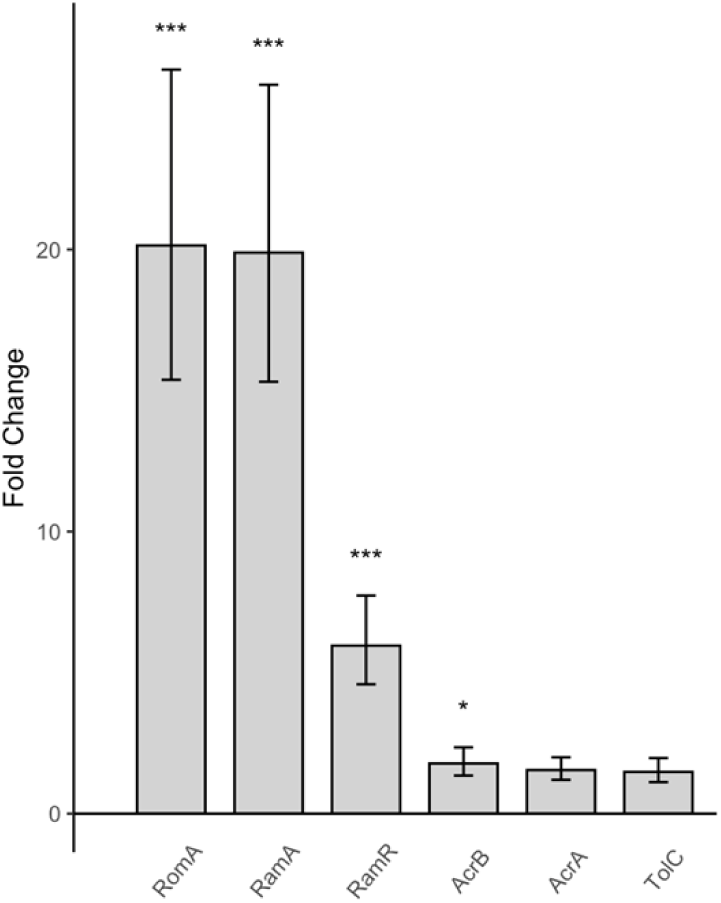
Differential Gene Expression between Strains with Functional *ramR* and Mutated *ramR*. Differential gene expression between strains with functionnal *ramR* (ESSO2 to ESSO6) and unfunctional *ramR* (ESSO7 and ESSO8) by quantitative PCR. Fold Changes calculated with ΔΔCt method and *p value* calculated with linear method. Significant *p value* was considered lower than 0.05 and represented by star.

### Global transcriptomic consequences of *ramR* mutations

Differential transcriptomic expression analysis between ESSO6 and ESSO7 was performed to identify genes impacted by the mutation of functional *ramR*. The detailed data are available in the supplementary data (S_transcriptomic analyses Appendix).

First, the transcription of 99 genes was significantly different between the two transcriptomes (Fig 4). Among these genes, 43 were found to be downregulated, and 56 were upregulated. The *ramAR* locus genes *ramA* (log2FC: 4.1) and *romA* (log2FC: 3.9) were upregulated, as was *ramR* (log2FC: 2.6) because of its autoregulation. Another transcriptional regulator, *acrR,* was significantly downregulated (log2FC:-1.09, *p value* adjusted: 0.04). Therefore, mutations in *ramR* impact the expression of two transcriptional regulators, *ramA* and *acrR,* which regulate numerous bacterial functions and could in turn be responsible for the dysregulation of other genes.

**Fig 4.**
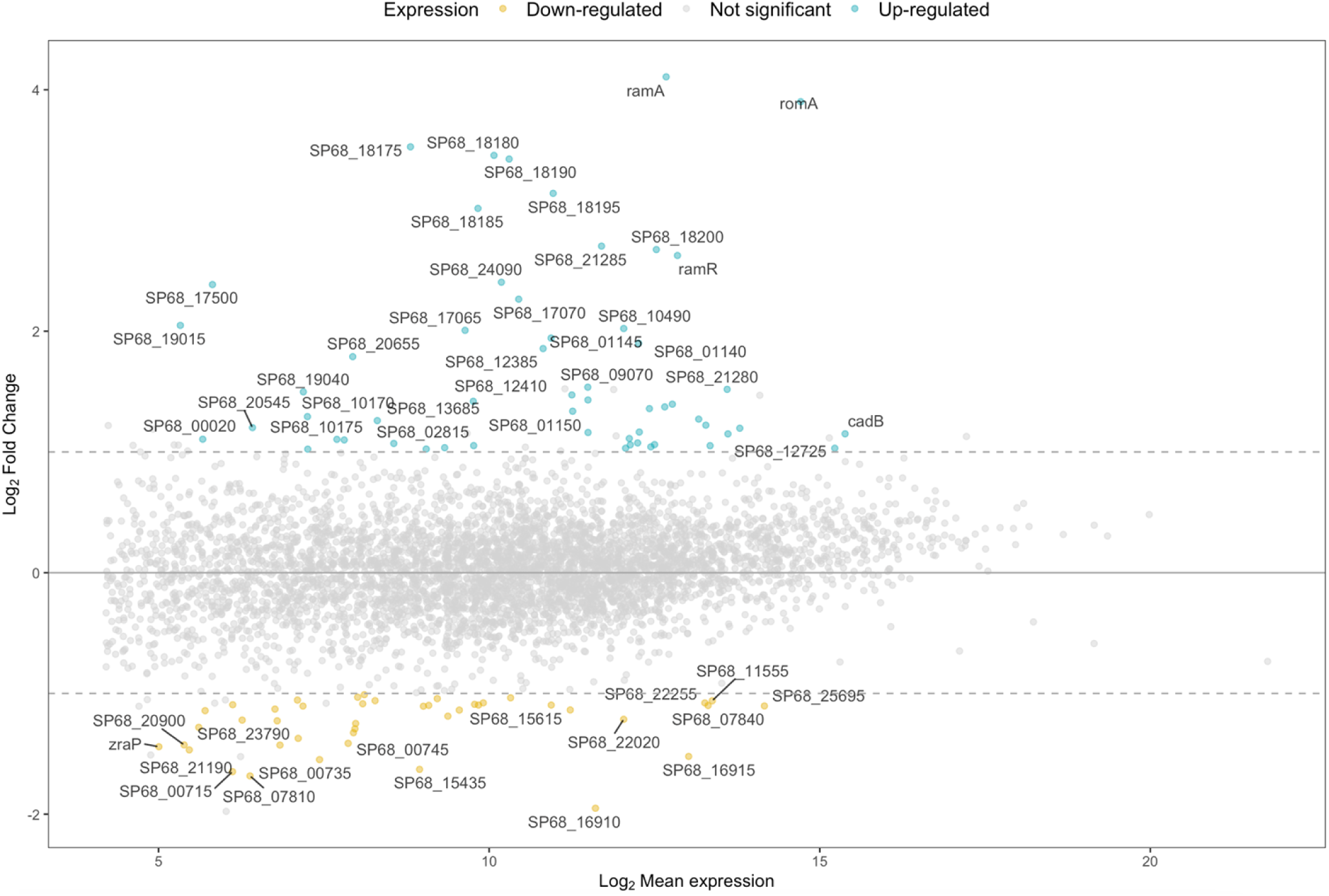
MA Plot of Differential Gene Expression between Isolates Unmutated and Mutated *ramR K. variicola*. MA plot of differential genes expression between not mutated *ramR K. variicola* (esso6) and *K. variicola* with mutation in *ramR* (esso7). Colored points represent genes with a significant *p value* adjusted with Benjamini & Hochberg method (*p value* adjusted <0.05), blue point for genes overexpressed and yellow points for genes under expressed. Dashed lines represent the threshold of Fold Change.

Several genes were significantly dysregulated with a log2FC of less than 1, including genes encoding the RND pumps *acrAB* (log2FC*_acrA_* 0.9, log2FC*_acrB_* 0.7) and *oqxAB* (log2FC*_oqxA_* 0.4, log2FC*_oqxB_* 0.6), *mla* operon (log2FC*_mlaC_*0.6, log2FC*_mlaD_* 0.7, log2FC*_mlaF_* 0.8) and the membrane porin OmpK37 (log2FC: 0.9), which all showed upregulation. Other membrane porins (OmpK35 and OmpC) did not show significant variation in expression. The upregulation of these pumps could explain the decreased susceptibility to chloramphenicol, tigecycline, quinolone and temocillin. The noncoding RNA *micF*, which regulates porin expression, was also upregulated, but not significantly, in ESSO7 (log2FC: 1.09, *p value* adjusted: 0.24).

Apart from the RND and porin transcripts, the *lsr* operon (log2FC*_lsrB_*: 1.47, log2FC*_lsrK_*: 1.06, log2FC*_lsrA_*: 0.75, log2FC*_lsrC_*: 0.69), encoding the autoinducer system/quorum sensing, was upregulated, and several genes encoding the phosphotransferase system, such as PTS sugar transporter subunit IIA (log2FC: 3.52), IIB (log2FC: 3.46), IIC (log2FC: 3.02) and IID (log2FC: 3.42), which are involved in capsule formation, were significantly upregulated. The gene *bhsA*, which encodes a stress resistance protein, was also upregulated (log2FC: 2.4). Conversely, the expression of many tRNAs were significantly downregulated, as were genes involved in aerobic/anaerobic metabolism, such as genes related to the PAA system (log2FC*_paaB_*:-1.65, log2FC*_paaJ_*:-1.43, and log2FC*_paaG_*:-1.37), which encode for phenylacetic catabolism and aerobic metabolism of aromatic compounds, and the *ybgE* gene (log2FC:-1.08), which encodes for a membrane protein implicated in aerobic metabolism. Genes encoding iron and hemin ABC transporters (SP68_09635, SP68_09630, SP68_09580 and SP68_09585) were also downregulated (S_transcriptomic analyses Appendix).

The genes *lipA* (log2FC: 0.67) and *lpxO* (log2FC: 0.83), which encode lipopolysaccharides, were slightly upregulated, as was the hemolysin *hha* transcriptional regulator (SP68_20685) (log2FC: 1.19), which is implicated in biofilm formation. The *mrk* operon (log2FC*_mrkD_*: 0.84, log2FC*_mrkF_*: 0.81, log2FC*_mrkC_*: 0.79) and another gene (SP68_01865) (log2FC: 0.99) associated with fimbria production and coding for type 3 fimbria were also moderately upregulated, while *fimC* (log2FC:-0.66), SP68_10605 (log2FC:-0.69) and SP68_10625 (log2FC:-1.08), other genes encoding fimbria proteins, were downregulated. Several genes involved in sugar metabolism were also upregulated (*treBC* for trehalose, *rsbD* for ribose, and several genes required for glucose metabolism). At final isolated genes were downregulated, including *zraP*, a zinc resistance sensor, and three efflux transporters (*kmrA*, *kpnF*, SP68_01265).

Finally, gene set enrichment analysis in the KEGG database was applied to the transcriptomic data (Fig 5). Seven metabolic pathways were significantly modified. Among them, three were downregulated (biosynthesis of cofactors, phenylalanine metabolism and tRNA biosynthesis), and four were upregulated (phosphotransferase system, TCA cycle, sucrose metabolism, and ribosome).

**Fig 5.**
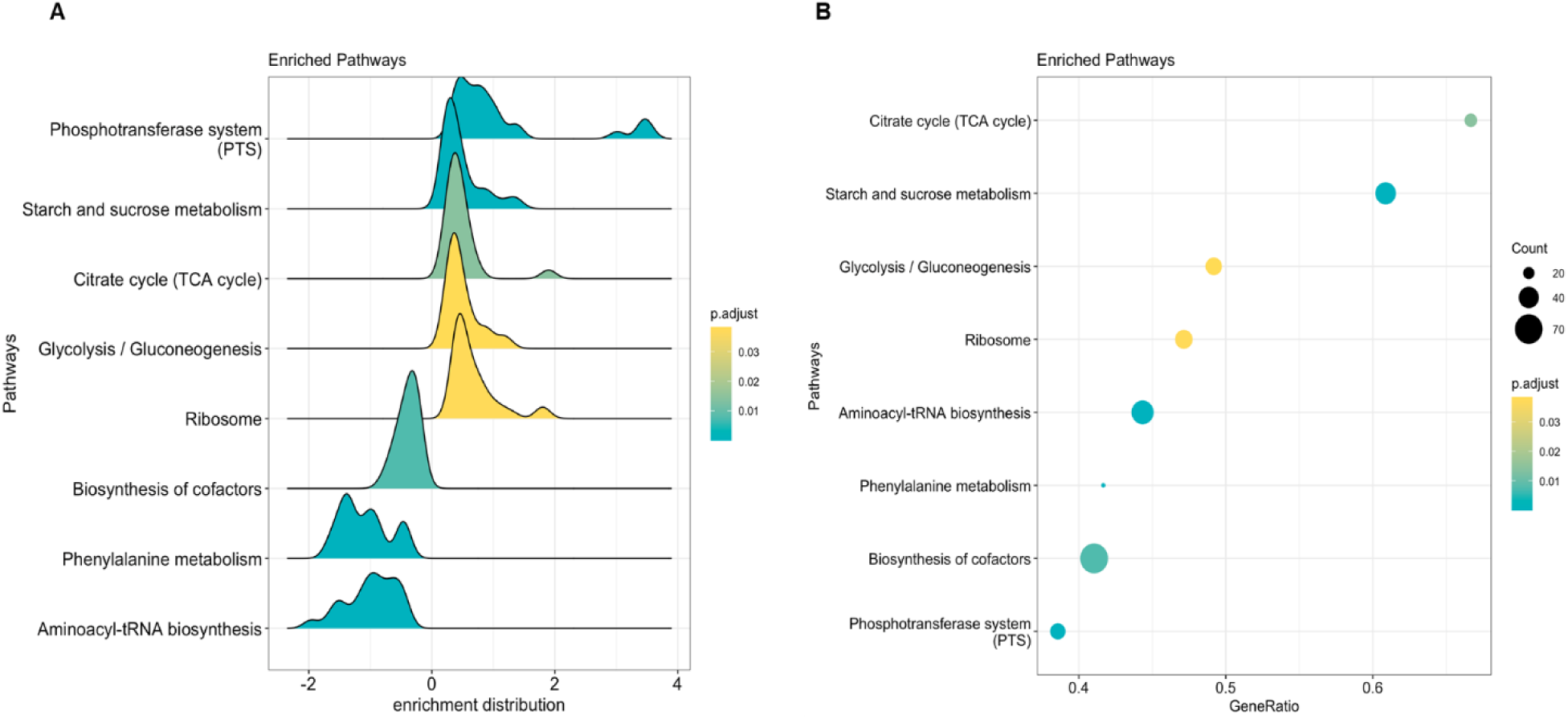
Gene Set Enrichment Analysis via KEGG Ontology. (A) Gene enrichment distribution and (B) gene ratio of enriched pathways between the ESSO6 and ESSO7 strains. *p.value* adjusted with the Benjamini & Hochberg method (R package “enrichplot”).

## Discussion

In this study, we followed the digestive carriage of KpSC populations though their excretion in the stool. In total, 38% (8/21) of patients hospitalized for more than 15 days in our ICU were colonized at least temporarily by a KpSC strain, and only 24% (5/21) excreted KpSC populations for several weeks. The colonization rate was approximately 10% (2/21) upon admission and increased to 33% (7/21) after three weeks of ICU stay. These results show an increase in the rate of KpSC colonization during the hospital stay, as highlighted in previous studies [4–7].

Among the five patients who showed KpSC colonization for several weeks, each had a unique persistent KpSC population until the end of hospitalization, except for one patient who experienced the disappearance of KpSC excretion after four weeks of hospitalization. SNP typing revealed a relatively stable KpSC population in each patient, with fewer than 12 SNPs between strains isolated from the same patient. SNP analysis of the core genome showed that all these ST strains were relatively close (median: 2 SNPs [min: 0, max: 4]) and were distinct from our local continuous ICU genomic ESBL-*Enterobacterales* surveillance [24]. The results from these five patients strongly support the hypothesis of persistent endogenic KpSC in the gut throughout a long hospitalization period.

In terms of antimicrobial resistance, two patients excreted isolates whose antimicrobial resistance profile changed during their ICU stay. The first patient had the *K. variicola* ST1791 strain, which acquired resistance genes likely contained in a plasmid and then showed rapid disappearance from the patient’s rectal samples. The second patient excreted long-term persistent *K. variicola* ST1563 digestive carriage isolates during an ICU stay of eight weeks. The *K. variicola* isolate collected at the end of hospitalization harbored mutations in an important RND pump regulator. Consequently, the antibiotic susceptibility profile was modified: resistance to tigecycline, quinolones and chloramphenicol appeared without the acquisition of resistance genes. Genetic exploration revealed a unique mutation in the *ramR* regulator that could have led to the RamR protein modification. RamR belongs to the TetR transcriptional regulator family and is well known to repress the expression of the *ramA* gene in *Enterobacterales* [25]. A unique mutation of *ramR,* which leads to an alteration of the function of the protein regulator, has already been described in other *Enterobacterales*, including *K. pneumoniae*, and results in a similar cross-resistance profile [26–28]. Furthermore, loss of the ability of *ramR* to regulate its targets affects not only antibiotic susceptibility but also biliary salt tolerance and has a pleiotropic effect on bacterial regulation [29,32]. However, unlike other studies, no MIC increase was observed for colistin in the strains with mutated *ramR*.

In a stress-free environment, the RamR protein binds to the *ramA* promoter, which is located next to the *ramR* locus, resulting in negative control of *ramA*. When a substrate molecule binds to the substrate domain site of the RamR protein, the *ramA* promoter is released, leading to its transcription [25,29]. RamA, a transcriptional regulator, can induce the expression of RND efflux pump genes, which could explain the decrease in antibiotic susceptibility [20,27]. The most broadly described RND pump in *Enterobacterales* is AcrAB, which is strongly linked to the *ramAR* locus [30,31]. In our case, a mutation in *ramR* resulted in an amino acid change, which seems to be in the substrate domain site of the protein and an amino acid directly implicated in hydrogen liaison with the substrate [32]. We validated the link between the *ramR* mutation and RND pump overexpression, which revealed increased expression of *ramA* in *the* mutated *ramR* strains compared with the unmutated strains. The AcrAB RND pump was moderately overexpressed, and *romA*, a gene strongly regulated by the *ramR* regulator, was also upregulated [33]. The involvement of the RND pump was confirmed with an RND pump inhibitor: in the presence of PAβN, the MICs of the mutated *ramR* strains decreased to those of the unmutated *ramR* strains.

Transcriptomic study of the *K. variicola* strains with a mutation in the *ramR* regulator revealed a change in the expression of *ramA* and *acrR*, two important transcriptional regulators with pleiotropic activities [32,34]. Changes in the expression of some RND pumps (*acrAB*, *oqxAB*, *mla* operons) could explain the decreased susceptibility to tigecycline, quinolones and chloramphenicol, as confirmed by the MICs. Interestingly, overexpression of OmpK37 porin does not reduce the MIC for β-lactam antibiotics as observed for other porins (OmpK35 and OmpK36) due to a lower affinity for these molecules, particularly for cefoxitin and cefotaxime [35]. Several genes involved in digestive colonization as *galET* and *waa* operon, which are involved in lipopolysaccharide formation, and *acrAB*, which is involved in anaerobic metabolism, were not found to be dysregulated in our transcriptomic analyses [13–15]. This could be explained by our study design that we did not compare the strains colonized from the patient’s gut at one moment but compare persistent strains in the digestive tract during an hospitalization course. However, we revealed other metabolic pathways that seem to be dysregulated because of the *ramR* mutation. Overexpression of genes related to the phosphotransferase system (PTS sugar transporter) and lipopolysaccharides (*lipA*, *lpxO*), which play roles in capsule and biofilm formation and virulence, was observed in the *ramR*-mutated strains. The transcriptomic changes observed could increase the mucoviscosity and biofilm formation of the strain and reduce its virulence [36]. However, further experiments are needed to confirm this hypothesis. Several other genetic changes were noted, including changes in genes related to quorum sensing (*lsr* operon), the stress response (*bhsA*), which is consistent with an increase in biofilm life and the downregulation of the phenylacetic catabolism and aerobic metabolism of aromatic compounds (PAA system, *ybgE*), and iron and hemin transport, which is consistent with a decrease in virulence [37]. These observations are consistent with the transcriptomic analysis of the RamA and RamR regulators carried out by De Majumdar on a laboratory-deleted *K. variicola* strain [34]. Indeed, the authors showed that the RamA regulator affected the expression of genes involved in biofilm formation, fimbria, LPS formation, virulence and sugar metabolism in *K. pneumoniae*. The RamR regulator that regulates the expression of RamA and RamR could indirectly be involved in the expression of these same bacterial functions. However, our analyses did not reveal variations in the nitrate reductase system (*nar* and *nir* operon). These differences could be explained by the type of strains studied in the different studies. In the De Majumdar study, the *ramR* gene was completely deleted, whereas in our study *ramR*, was only mutated and could still show activity at some of these targets.

This study is the first report, to our knowledge, of *ramR* mutations in *K. variicola* strains isolated during a longitudinal clinical follow-up, during which we were able to observe KpSC isolates regardless of the AMR profile present in the digestive flora of patients with a prolonged ICU stay. Consequently, we observed the adaptations of these isolates throughout the hospitalization stay. The *ramR* mutation and subsequent phenotypic changes may be a key to bacterial persistence, possibly by modifying its capacity to form biofilms and improving its tolerance to biliary and antibiotic pressure.

## Materials and methods

### Collection of *K. pneumoniae* species complex populations

#### Patient recruitment and sample collection

This longitudinal study took place between December 2018 and May 2019 in two adult ICUs, one medical (22 beds) and one surgical (35 beds), at Caen University Hospital, a 1410-bed teaching hospital in Normandy, France. The inclusion process was performed from December 2018 to February 2019, and the sampling of included patients continued until May 2019. The samples collected were rectal swabs used for *Enterobacterales* ESBL or/and CP detection in ICUs following local practices, including weekly sampling of each hospitalized ICU patient throughout the hospital stay. This protocol was in accordance with national recommendations to adapt presumptive antibiotic treatment for infection and establish preventive hygiene measures for the patient, as recommended by the “Société Française d’Anesthésie et de Réanimation” [38]. The swabs used were transport swabs (Copan, Brescia, Italy). The inclusion criteria were as follows: patients who were at least 18 years old, had no major comorbidities, and had no history of hospitalization or antibiotic treatment for the three months prior to hospitalization. Furthermore, the length of ICU stay had to be for a continuous period of more than 15 days and the first rectal swabs had to be taken within one week after admission. Further, at least three swabs needed to be collected during the hospital stay.

#### Patient data

For the patients included, demographic data were collected from paper and electronic medical records (Usv2-Crossway, McKesson, Irving, TX, USA). The information obtained included age, sex, all treatments received, hospitalization duration, outcomes during the hospital stay, medical history and post-ICU orientation. Laboratory software (TDNexLabs, Technidata, Montbonnot-Saint-Martin, France) was used to collect microbiology laboratory sample information.

#### *Klebsiella* detection and isolates

The collected swabs were cultivated for 24 h at 35± 2 °C in Luria–Bertani (LB) broth supplemented with amoxicillin (10 mg/L) and vancomycin (16 mg/L) to reduce the bacterial charge without affecting KpSC growth. Next*, Klebsiella* isolates were obtained from the LB cultures via a two-step protocol: (i) ZKIR PCR on all samples and (ii) culture on Simmons-citrate-agar-inositol (SCAI) selective media, as previously described [39,40].

Total bacterial DNA was extracted from LB broth cell pellets by InstaGene Matrix (Bio-Rad, Marnes-la-Coquette, France), and real-time ZKIR PCR was performed using Rotor-Gene Q (Qiagen, Hilden, Germany) and the Takyon ROX Probe Mastermix kit (Eurogentec, Seraing, Belgium). Broth samples with positive PCR were plated on SCAI media after serial dilution (10^-4^, 10^-5^ and 10^-6^) to obtain nonconfluent cultures [40]. After 48 hours of incubation at 35 ± 2 °C, identification with MALDI-TOF technology (Bruker Daltonik, Bremen, Germany) was performed on characteristic colonies. Finally, five distinct KpSC colonies were isolated from each positive sample and were preserved in vials of BHI supplemented with glycerol at-80 °C. The control strains used for this study were *K. pneumoniae* ATCC 700603 and *K. variicola* ATCC BAA-830, which were conserved under the same conditions as the *Klebsiella* collection.

All the patients with long-term KpSC carriage were anonymized under the acronyms BOMA, GUMA, ESSO, ETJP, and OZAR.

### Biocide survey

#### Antimicrobial susceptibility testing

A large screening of antimicrobial susceptibility profiles, with a panel of 35 antimicrobial agents (Bio-Rad, Marnes-la-Coquette, France), was performed using the disk diffusion method on MH agar (BD, Heidelberg, Germany) for each *Klebsiella* isolate. The antibiotics used are listed in the supplementary S_material and methods Appendix. Decreased susceptibility (more than 5 mm between two strains) during longitudinal surveillance was confirmed by MICs in broth microdilution in 96-well plates measured in triplicate. The two antibiogram methods were performed and interpreted as recommended by the Comite de l’antibiogramme de la Société Francaise de Microbiologie/European Committee on Antimicrobial Susceptibility Testing 2020 guidelines (CA-SFM/EUCAST: https://www.sfm-microbiologie.org/).

#### Antiseptic susceptibility testing

Antiseptic susceptibilities were assessed in MH (BD, Heidelberg, Germany) broth microdilutions in 96-well plates for three antiseptics commonly used at hospitals: alcoholic chlorhexidine 2% (Gilbert laboratories®, Hérouville-Saint-Clair, France), povidone-iodine 10% (Betadine®, Meda Pharma, Paris, France) and DDAC (Anios®, Lezennes, France), a quaternary ammonium commonly found in hospital surface disinfectants. Supplementary data are provided in the supplementary S_material and methods Appendix. The plates were inoculated with a 1/10 dilution of bacterial suspensions calibrated at McFarland 0.5 and incubated at 35 ± 2 °C for 24 hours. For each molecule, the first three wells with no visible growth were plated on TS and incubated for 24 hours at 35 ± 2 °C to determine the minimal bactericidal concentration (MBC).

### Genomic analyses

Whole-genome sequencing and analyses were carried out on one or two isolates collected per week based on their antimicrobial susceptibility profile changes. In total, 24 isolates were sequenced for five KpSC-excreting long-stay patients (S1 Appendix).

#### DNA extraction and short-read whole-genome sequencing

DNA of KpSC isolates was extracted from cells incubated overnight in LB broth and sequenced at the Plateforme de Microbiologie Mutualisée (P2M-Institut Pasteur, Paris, France). DNA was extracted with a MagNAPur 96 system (Roche Applied Science, Mannheim, Germany). Whole-genome sequencing was performed with an Illumina NextSeq 500 instrument (San Diego, CA, USA) using a Nextera XT DNA Library Preparation Kit, and 150 bp paired-end reads were generated. The paired-end reads were subjected to quality selection with fqCleaner from AlienTrimmer [41] and were assembled *de novo* using SPAdes V3.12.0 [42]. The quality of the genome assembly was assessed using Quast software [43].

#### Bioinformatic analyses

The identification, multilocus sequence typing (MLST) characterization, and virulence and resistance genes of each sequenced isolate were identified with Kleborate 0.4.0 [44,45]. As only *K. pneumoniae* and *K. variicola* were identified in our study, a core genome MLST (cgMLST) was performed with ChewBBACA 2.8.5 throughout the cgMLST gene database (https://www.cgmlst.org/ncs/schema/2187931/) [46]. The distance matrix generated was compared with that of our local database, implemented since 2015, as part of the bacterial resistance genomic survey. Neighbor-joining trees were generated with GrapeTree and iTOL tools. Single nucleotide polymorphism (SNP) analysis of all *Klebsiella* genomes isolated from the same patient was performed with Snippy 4.6.0 [47], and the results were visualized with IGV software [48]. For the mapping process, the genome of the first isolate found from each patient was used as a reference. Coding DNA sequences (CDSs) were identified and annotated with Prokka 1.14.5 [49]. A pairwise core-genome SNP analysis was performed between isolates.

Virulome, resistome, cgMLST and SNP analyses of all isolates from each patient were compared to observe the evolution of genomic dynamics and identify adaptation mechanisms.

### Transcriptomic analysis of the ESSO *K. variicola* isolates

#### RNA extraction

ESSO2 to ESSO8 were cultivated under the same conditions. Overnight cultures in LB broth were inoculated and incubated at 35 ± 2 °C with shaking. Next, 100 µL of the overnight culture was added to 10 mL of LB broth, incubated with shaking at 35±2 °C until the end of the exponential phase (±4.5 hours), and then centrifuged. RNA extractions were performed on bacterial pellets.

Total RNA extraction was performed with the Direct-Zol RNA mini-prep kit (Zymo Research, Irvine, CA, USA), and residual DNA was removed with the TURBO DNA-free Kit (Thermo Fisher Scientific, Wilmington, MA, USA), according to the manufacturer’s protocol. The extracts were quantified with a NanoDrop One spectrophotometer (Thermo Scientific, Wilmington, MA, USA) and TapeStation RNA ScreenTape (Agilent Technologies, Santa Clara, CA, USA).

#### RNA sequencing and analyses

The transcriptomes of ESSO6 (the last isolate without the *ramR* mutation) and ESSO7 (the first isolate with the *ramR* mutation) were explored. Triplicates of RNA extracts obtained as previously described were sequenced with an Illumina HiSeq 4000 sequencer using Total RNA Prep with Ribo-Zero Plus Kit at the iGE3 Genomics Platform (University of Geneva, Swiss), generating single reads of 100 nucleotides with ribosomal RNA depletion.

The RNA sequence quality check was performed with a FastQC analyzer (version 0.11.9). RNA sequences were then mapped to the reference genome of *K. variicola* ATCC BAA-830 (RefSeq sequence: NZ_CP010523.2) with STAR mapper (version 2.7.9a) [50]. A gene count table was obtained with HTseq-count (version 1.99.2). Normalization and differential expression analysis of RNA counts were performed with the R package DESeq2 (version 1.34.0) [51]. The *p values* were calculated by the Wald test and corrected with the Benjamini and Hochberg method. Gene transcripts were compared for those with *p values* ≤ 0.05 and log2 fold changes (log2 FC) >1. The data are available in the S_transcriptomic_analyses Appendix. Next, Kyoto Encyclopedia of Genes and Genomes (KEGG) enrichment was performed to identify the potential metabolic pathways impacted.

#### Gene expression by RNA quantification

RNA extracts were reverse transcribed into cDNA using an iScript™ Select cDNA Synthesis Kit (Bio-Rad, Marnes-la-Coquette, France) following the manufacturer’s instructions. RNA and cDNA extracts were stored at-80 °C.

Gene-specific primers for *ramA, ramR, romA, acrA, acrB, tolC* and *gyrA* were designed with Primer3 (https://www.ncbi.nlm.nih.gov/tools/primer-blast/) using the ESSO2 strain as the reference (S_Materials and Methods Appendix). Fold changes were calculated with the ΔΔCT method using *gyrA* as a housekeeping gene [52]. RNA expression between isolates with wild-type *ramR* (ESSO2 to ESSO6) and isolates with mutated ramR (ESSO7 and ESSO8) was compared via Student’s test. The results with a *p value* lower than 0.05 were considered statistically significant.

### *RamAR* locus investigations

The *ramAR* locus was identified as a potential gene involved in the adaptation of the KpSC isolates. Therefore, an investigation of the impact of this gene mutation was performed.

#### Phenylalanine-arginine beta-naphthylamide (PAβN) test

The MICs of the broth microdilutions to antibiotics (chloramphenicol, tigecycline, ciprofloxacin, and temocillin) in the presence of PAβN (Sigma‒Aldrich, Saint-Louis, MO, USA), an RND pump inhibitor, were determined to validate the hypothetical link between *ramAR* locus mutation, pump overexpression and decreased antibiotic susceptibility. The PAβN concentration was fixed at 20 mg/L [53].

#### Gene complementation

Complementation with the functional *ramR* gene was performed on isolates carrying a mutated *ramR* gene. The bacterial DNA of the clinical strain containing functional *ramR* (ESSO2 strain) was extracted, and then, *ramR* was amplified by PCR using specific *ramR* primers (S_Materials and Methods Appendix). The amplified *ramR* gene was inserted into the pBAD202 vector (pBAD202 Directional TOPO™ Expression Kit – Invitrogen, Carlsbad, CA, USA), transformed in One Shot™ TOP10 chemically competent *E. coli* (Thermo Fisher Scientific, Wilmington, MA, USA) and purified as described by the manufacturer. Complementation of the targeted strains (ESSO7 and ESSO8) with pBAD202 containing a *ramR* insert was performed by electroporation. Complemented strains were selected under kanamycin (40 mg/L) pressure. Complementation with empty pBAD202 was performed to confirm the absence of an effect on bacterial strain fitness and antimicrobial susceptibility profile.

#### Phenotypic analysis: Growth curves in a microplate reader

Fitness tests and growth curve generation were performed with a Tecan Infinite 200 Pro microplate reader (Tecan, Männedorf, Switzerland) on ESSO2 to ESSO8 and on the complemented ESSO strains. Growth curves were generated in triplicate in 96-well plates in LB broth, starting with a bacterial inoculum of 10^7^ CFU/mL obtained from an overnight culture. The plates were shaken every 10 minutes, and the optical density (OD) was read at 600 nm every 30 minutes for 15 hours. Second, growth curves of the ESSO strains were generated under different stress conditions to evaluate the adaptability of the isolates to the following: biliary salts (40 and 20 mg/mL), hyperosmotic stress (NaCl at 30 and 15 mg/mL), pH (4.5 to 5.5), temperature (37 °C and 22 °C), starvation (M9 medium), erythromycin (256 to 64 mg/L), and oxidative stress (H_2_O_2_ at 1 and 0.25 mM). The generation time, inflection time and curve regression were calculated with the “Growthcurver” R package and compared [54].

### Statistical analysis and graphic representation

Statistical analysis, graphic representation and statistical tests were performed with RStudio software (version 1.2.1335). Student’s tests were performed to compare the qPCR results (isolates with mutated *ramR* versus isolates without mutated *ramR*). Student’s tests were performed on all growth curve parameters to compare isolates with or without the mutated *ramR* gene and to compare complemented isolates with or without functional *ramR*.

## Acknowledgements

We are grateful to Agathe CAPITAINE, Michel AUZOU, Mamadou GODET and Sebastien GALOPIN for their technical support. We are thankful to the ICU supervisors for their contribution to this project, in particular Frederic ETHUIN.

## Ethics approval

This study received ethical approval from the Committee for the Protection of Persons of the CHU of Caen (ID-RCB: 2020-A00479-30).

## Author contributions

BL and F Gravey participated in the methodology, investigation, formal analyses, data curation and writing. F Gravey, SLH and CI participated in the conceptualization, methodology and supervision. AG, F. Guerin and MA participated in the investigation. SB, JCG, FE, DDC and CG participated in the methodology and visualization. All the authors reviewed the manuscript.

## Data availability

The genome assemblies and SRAs of the RNA-seq sequences are available in BioProject (PRJNA1053322).

## Supporting information

Supplementary data are available in:

S1_S2_S5_S6_tables_figures Appendix

S3_kleborate Appendix

S4_all_diameters Appendix

S7_growthcurver_data Appendix

Supplementary S_transcriptomic_analyses Appendix

Supplementary materials and methods are available in:

Supplementary S_material and methods Appendix

